# Prediction of multiple drug resistant pulmonary tuberculosis against drug sensitive pulmonary tuberculosis by CT nodular consolidation sign

**DOI:** 10.1101/833954

**Authors:** Xi-Ling Huang, Aliaksandr Skrahin, Pu-Xuan Lu, Sofia Alexandru, Valeriu Crudu, Andrei Astrovko, Alena Skrahina, Jessica Taaffe, Michael Harris, Alyssa Long, Kurt Wollenberg, Eric Engle, Darrell E. Hurt, Irada Akhundova, Sharafat Ismayilov, Elcan Mammadbayov, Hagigat Gadirova, Rafik Abuzarov, Mehriban Seyfaddinova, Zaza Avaliani, Sergo Vashakidze, Natalia Shubladze, Ucha Nanava, Irina Strambu, Dragos Zaharia, Alexandru Muntean, Eugenia Ghita, Miron Bogdan, Roxana Mindru, Victor Spinu, Alexandra Sora, Catalina Ene, Eugene Sergueev, Valery Kirichenko, Vladzimir Lapitski, Eduard Snezhko, Vassili Kovalev, Alexander Tuzikov, Andrei Gabrielian, Alex Rosenthal, Michael Tartakovsky, Yi Xiang J Wang

**Affiliations:** Department of Imaging and Interventional Radiology, Faculty of Medicine, The Chinese University of Hong Kong, Hong Kong SAR, China; Belarusian State Medical University, Minsk, Republic of Belarus; Republican Scientific and Practical Centre of Pulmonology and Tuberculosis, Ministry of Health, Minsk, Republic of Belarus; Shenzhen Center for Chronic Disease Control, Shenzhen, Guangdong Province, China; Phthysiopneumology Institute, Ministry of Health, Chisinau, Republic of Moldova; Office of Cyber Infrastructure and Computational Biology, National Institute of Allergy and Infectious Diseases, National Institutes of Health, Department of Health and Human Services, Bethesda, Maryland, USA; Scientific Research Institute of Lung Diseases, Ministry of Health, Baku, Republic of Azerbaijan; The National Center for Tuberculosis and Lung Diseases, Tbilisi, Republic of Georgia; Marius Nasta Pneumophtisiology Institute, Ministry of Health, Bucharest, Romania; United Institute of Informatics Problems, National Academy of Sciences of Belarus, Minsk, Republic of Belarus

## Abstract

Multidrug-resistant tuberculosis (mdrtb) refers to TB infection resistant to at least two most powerful anti-TB drugs, isoniazid and rifampincin. It has been estimated that globally 3.5% (which can be much higher in some regions) of newly diagnosed TB patients, and 20.5% of previously treated patients had mdrtb. Extensively drug-resistant TB (xdrtb) has resistance to rifampin and isoniazid, as well as to any member of the quinolone family and at least one of the second line injectable drugs: kanamycin, amikacin and capreomycin. xdrtb accounts for 4-20% of mdrtb. Early detection and targeted treatment are priorities for mdrtb/xdrtb control. The suspicion of mdr/xdr -pulmonary TB (mdrptb or xdrptb) by chest imaging shall suggest intensive diagnostic testing for mdrptb/xdrptb. We hypothesize that multiple nodular consolidation (NC) may serve one of the differentiators for separating dsptb vs mdrptb/xdrptb cases. For this study, mdrptb cases (n=310) and XDR-PTB cases (⋂=I58) were from the NIAID TB Portals Program (TBPP) <https://tbportals.niaid.nih.gov>. Drug sensitive pulmonary TB (dsptb) cases were from the TBPP collection (n=112) as well as the Shenzhen Center for Chronic Disease Control (n=111), Shenzhen, China, and we excluded patients with HIV(+) status. Our study shows NC, particularly multiple NCs, is more common in mdrptb than in dsptb, and more common in xdrptb than in mdrptb. For example, 2.24% of dsptb patients, 13.23% of mdrptb patients, and 20.89% of xdrptb patients, respectively, have NCs with diameter >= 10mm equal or more than 2 in number.

## Introduction

Multidrug-resistant tuberculosis (MDR-TB) refers to TB infection resistant to at least two most powerful anti-TB drugs, isoniazid and rifampincin. Extensively drug-resistant TB (XDR-TB) has resistance to rifampin and isoniazid, as well as to any member of the quinolone family and at least one of the second line injectable drugs: kanamycin, amikacin and capreomycin. XDR-TB accounts for 4–20% of MDR-TBs (1, 2). It has been estimated that globally 3.5% (which can be much higher in some regions) of newly diagnosed TB patients, and 20.5% of previously treated patients had MDR-TB (3). Early detection and targeted treatment of MDR-TB are priorities for MDR-TB control (4). Though geno- and phenotypic tests are available to detect *Mycobacterium tuberculosis (M.tb)* strains and their susceptibility/resistance to anti-TB drugs, delay of MDR-TB diagnosis and treatment is common. These tests can still produces inconclusive results and may not be available in resource-poor localities. The suspicion of MDR/XDR-pulmonary TB (mdrptb or xdrptb) by chest imaging shall suggest intensive diagnostic testing for MDR/XDR-TB.

During the course of our study reviewing CT images of drug-sensitive pulmonary TB (dsptb), mdrptb, and xdrptb, we conceived the possibility that multiple nodular consolidation may serve one of the differentiators for separating dsptb vs mdrptb/xdrptb cases. This possibility has also been recently suggested by other authors, though not specifically and not quantitatively (5, 6). This study meticulously quantified the number and size of NCs in a large collection of PTB database with CT images, and suggested some statistics that may help clinical decision making during PTB management.

## Materials and Methods

The MDR-PTB cases and XDR-PTB cases were from the NIAID TB Portals Program (TBPP) <https://tbportals.niaid.nih.gov> (7). dsptb cases were from base the TBPP collection as well as the Shenzhen Center for Chronic Disease Control, Shenzhen, China (table-1). We excluded patients with HIV(+) status, as patients with compromised immunofunction have been noted to have different ptb presentation (8). The category of being dsptb, mdrptb, or xdrptb was based on either (1) sputum cultured *M.tb* drug sensitivity testing, or (2) *M.tb* molecular testing results, (3) clinically diagnosed, i. e. favorable response and outcome following standard treatment regimen for dsptb (table-2).

**Table-1.** source of the TPB cases analyzed in this study.

| country | dsptb | mdrptb | xdrptb |
| --- | --- | --- | --- |
| | n=, (mean age±sd yrs) | n=, (mean age±sd yrs) | n=, (mean age±sd yrs) |
| Belarus (n=432) | 94, (46.9±17.2) | 214, 41.8±14.4 | 124, 39.3±12.9 |
| China (n=111) | 111, (33.8±11.9) | 0 | 0 |
| Romania (n=97) | 3, (29.7±15.1) | 67, (41.7±13.4) | 27, (42.9±11.7) |
| Azerbaijan (n=31) | 13, (36.7±16.6) | 15, (40.0±14.1) | 3, (28.3±4.0) |
| Moldova (n=17) | 1, (43) | 12, (35.7±15.2) | 4, (43.8±14.1) |
| Georgia (n=3) | 1, (20) | 2, (24.5±12.0) | 0 |
| total | 223, (39.4±15.9) | 310, (41.4±14.2) | 158, (39.8±12.7) |

**Table-2.** diagnostic method for assessing anti-TB drug sensitivity status.

| diagnostic method | dsptb (other than China) | dsptb (China) | mdrptb | xdrptb |
| --- | --- | --- | --- | --- |
| *M.tb* culture DST | 72 | 64 | 309 | 157 |
| *M.tb* molecular test | 1 | 12 | 1 | 1 |
| clinically diagnosed | 39 | 35 | 0 | 0 |

All the CT images were ready by at least one reader (XLH), a radiology trainee with two years’ experience in reading PTB imaging, and an experienced radiologist. The final reading was based on consensus, with input from a third radiologist when disagreement occurs. The current report focus on the CT feature of nodular consolidation (NC). For all NCs with diameter ≥6 mm, the number was counted for each patient; the diameter for each NC was measured on axial CT images showing the largest size and the mean of two perpendicular lengths was taken as the size of the NC. Some examples of the nodules are shown in Fig-1, and the measurement of a NC is illustrated in Fig-2.

**Fig-1.**
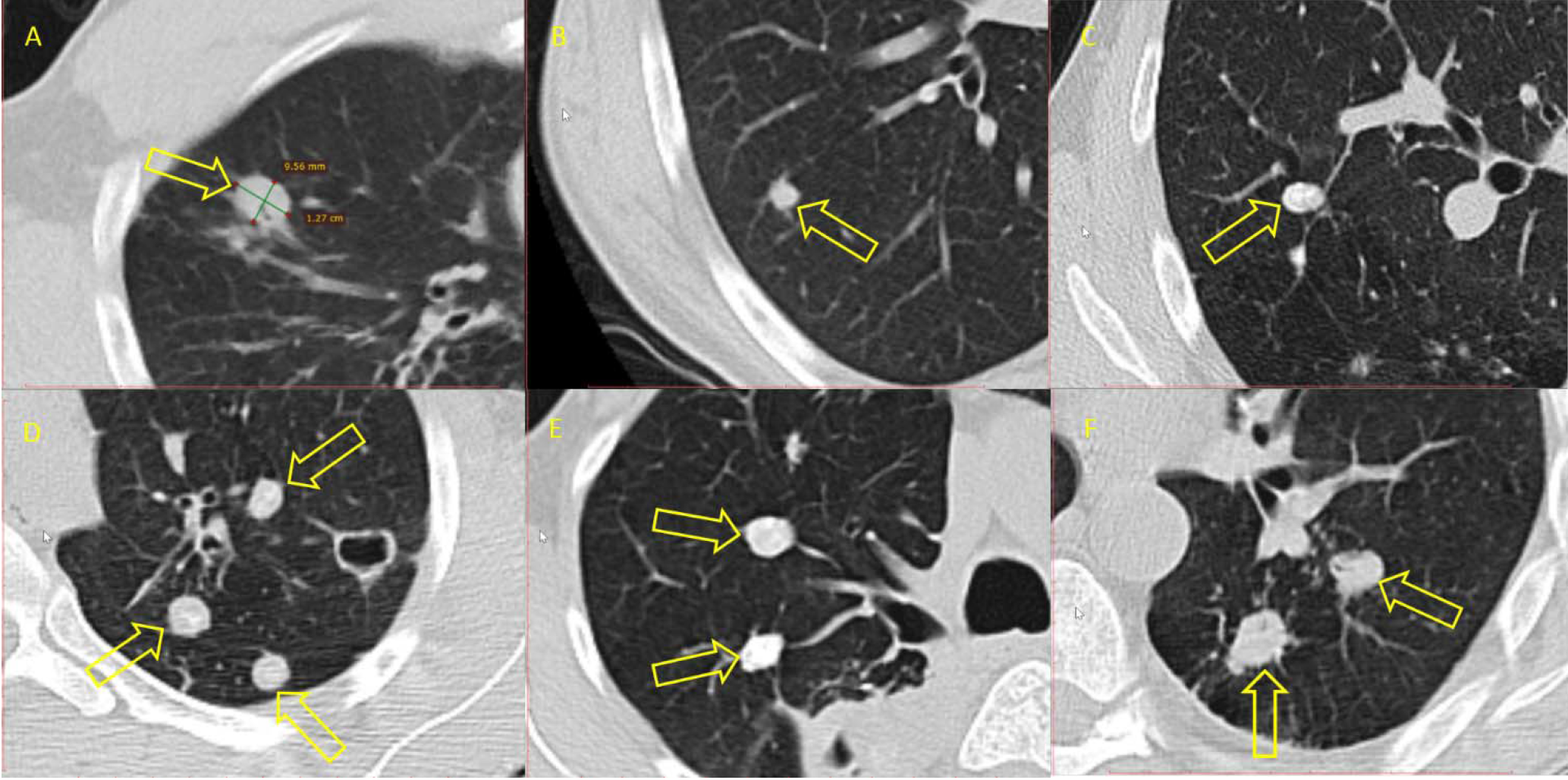
Examples of nodular consolidation (NC). A and B are from dsptb cases; C and D are from mdrptb cases; and E and F are from xdrptb cases.

**Fig-2.**
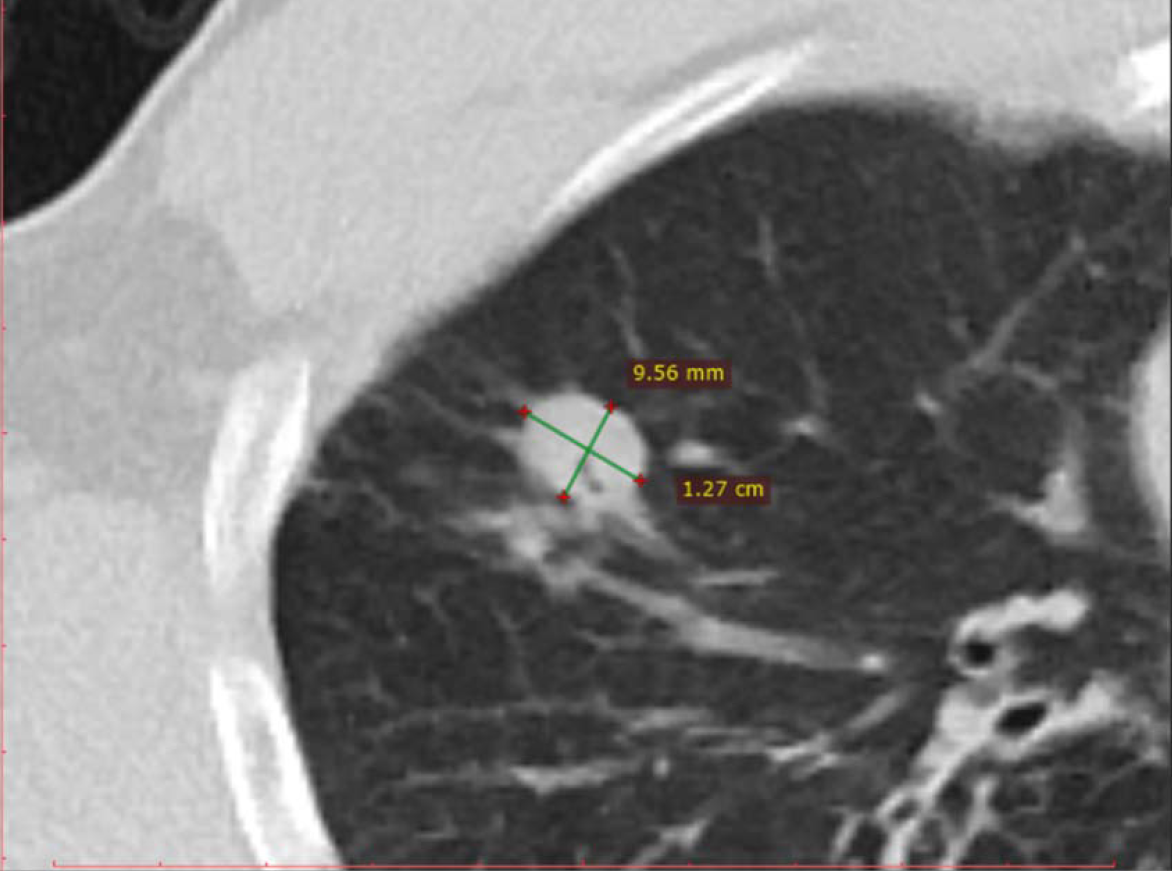
An example of measurement of nodular consolidation (NC)’s diameter. The diameter for each NC was measured on axial CT images showing the largest size and the mean of two perpendicular lengths was taken as the size of the NC.

## Results

Results of NC analysis for the study cohorts are shown in Fig-3, and examples are given in table-3. Moreover, for inter-regional comparison, our preliminary analysis shows there was no apparent NC prevalence difference between the Eastern Europe/Caucasus dsptb patients and Chinese dsptb patients (Fig-4).

**Fig-3.**
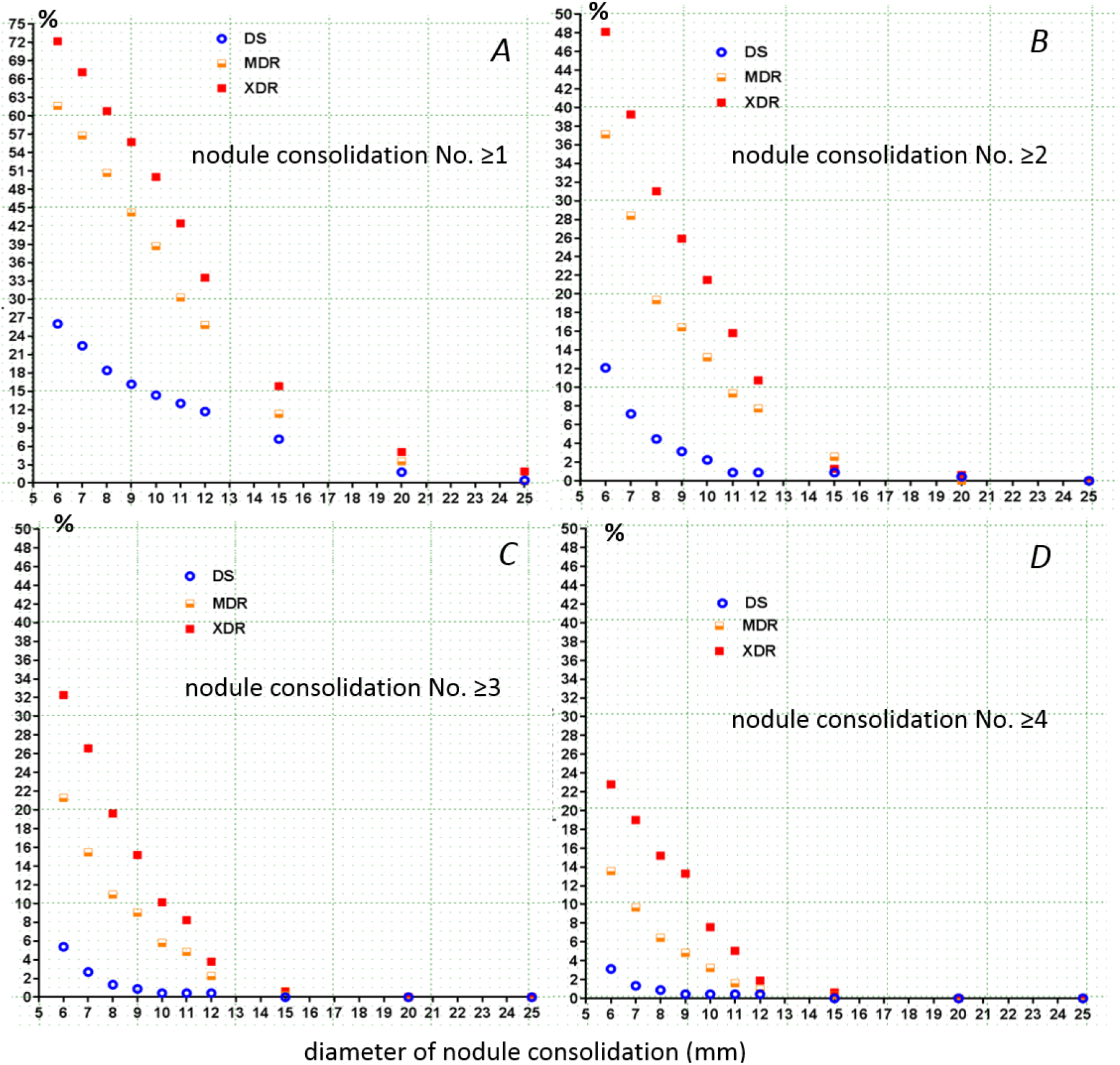
Prevalence of nodular consolidation (NC) of certain sizes in dsptb (DS), mdrptb (MDR), and xdrptb (XDR) patients. X-axis, NC diameter in mm. Y-axis, percentage prevalence of NC number of ≥1 (A), ≥2(B), ≤1(C), and ≤1 (D). For example, it can be seen that A shows approximately 72% of xdrptb cases have at least one NC with diameter equal or larger than 6mm; C show approximately 1% of dsptb cases have at least three NCs with diameter equal or larger than 9mm.

**Fig-4.**
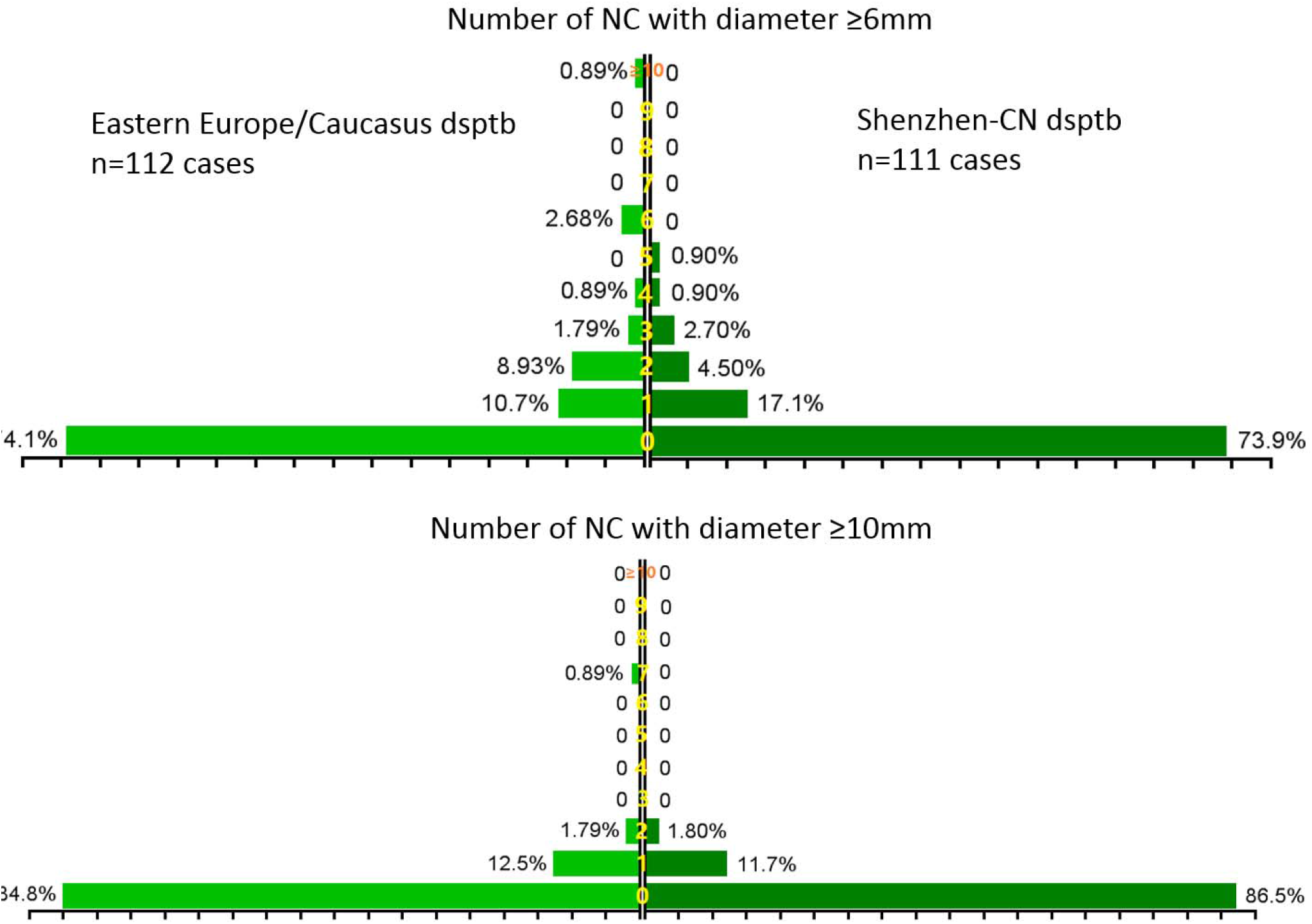
Prevalence of nodular consolidation (NC) of ≥6 mm or ≥10 diameter among dsptb patients from Eastern Europe/Caucasus and from China (Shenzhen). Transverse-axis: prevalence of NC; longitudinal-axis: number of NCs in 0, 1, 2, 3, 4, 5, 6, 7, 8, 9, and ≥ 10 respectively. This graph shows there is no apparent NC prevalence difference between the Eastern Europe/Caucasus dsptb patients and Chinese dsptb patients.

**Table-3.**
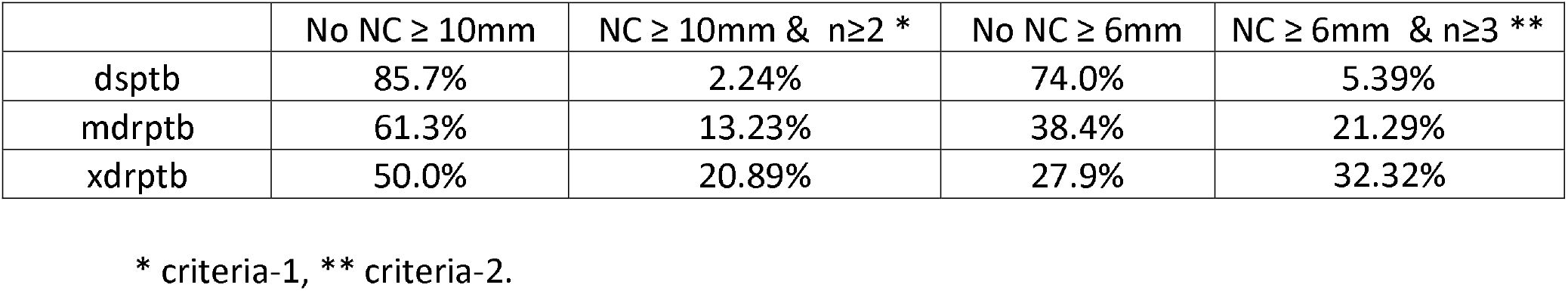
Selected criteria for predicting the probability based on nodular consolidation (NC) size and number.

| | No NC ≥ 10mm | NC ≥ 10mm & n≥2 * | No NC ≥ 6mm | NC ≥ 6mm & n≥3 ** |
| --- | --- | --- | --- | --- |
| dsptb | 85.7% | 2.24% | 74.0% | 5.39% |
| mdrptb | 61.3% | 13.23% | 38.4% | 21.29% |
| xdrptb | 50.0% | 20.89% | 27.9% | 32.32% |

\* criteria-1, \*\* criteria-2.

If the mdrptb prevalence in a specific population is 5% for new ptb patients, then for 1000 new patients (950 cases of dsptb and 50 cases of mdrptb):

1. 950*2.24% =21.28 patients will have NCs with diameter ≥10mm equal or more than 2 in number (criteria-1).
2. since these 1000 new patients will have 50 cases of mdrptb, these cases will have 50*13.23%=6.615 cases meeting criteria-1.
3. if we see a new patient meeting criteria-1, then probability of 6.615/(21.28+6.615)= **23.7%** of this patient will be a mdrptb.

--------

1. 950*5.39%= 51.205 patients will have NC with diameter ≥6mm and equal or more than 3 in number (criteria-2)
2. since these 100 new patients will have 50 cases of mdrptb, these cases will have 50*21.29%=10.645 cases meeting criteria-2.
3. if we see a new patient meeting criteria-2, then probability of 10.645/(50.1205 +10.645)= **17.2%** of this patient will be a mdrptb.

If the mdrptb prevalence in specific population is 20% for re-treatment ptb patients, then for 1000 retreatment patients (800 cases of dsptb and 200 cases of mdrptb) and if the CT feature of new dsptb and re-treatment mdrptb patients are the same, then:

1. 800*2.24% =17.92 dsptb patients will have NC with diameter ≥10mm and equal or more than 2 in number (criteria-1)
2. since these 1000 new patients will have 200 cases of mdrptb, these cases will have 200*13.23%=26.46 cases meeting criteria-1.
3. If we see a re-treatment patient meeting criteria-1, then a probability of 26.46/(17.92+26.46)= **59.6%** of this patient will be a mdrptb.

--------

1. 80*5.39% =4.312 dsptb patients will have NC with diameter ≥6mm and equal or more than 3 in number (* criteria-2)
2. since these 1000 re-treatment patients will have 20 cases of mdrptb, these cases will have 20*21.29%= 4.258 cases meeting criteria-2.
3. if we see a re-treatment patient meeting criteria-2, then a probability of 4.258/(4.312+4.258)= **49.7%** of this patient will be a mdrptb.

A high probability of mdrptb should trigger intensive diagnostic testing such as sputum microbiology culture drug sensitivity or GeneXpert testing (since the overall prevalence of xdrptb is very low, unless distinct signs for xdrptb is found. Here we do not offer examples for estimating xdrptb probability).

## Discussion

In a review published in early 2018, we summarized the radiological features for mdrptb (8). Till the end of 2017, the importance of multiple NCs as an imaging suggesting mdrptb was not recognized. Differentiation of dsptb and mdr/xdrptb has generally been considered impossible. For the computer assisted analysis, it was noted that no solution has sufficient prediction accuracy (10–14). Our study suggests a combination of NC number and size can be used to predict the probability of mdrptb, taking into account of the local prevalence of mdrptb. It can be expected that the quantification of NC number and size can be facilitated by computer aided detection. We are actively working to (1) increase the sample size of dsptb, (2) analyze and incorporate the mdrptb cases in Shenzhen Center for Chronic Disease Control into our mdrptb case pool, (3) incorporate other imaging features such as cavitary lesion and TB lesion lobular distribution etc into the prediction model. It has been suggested that multiple cavities, particularly those with thick wall, are associated with mdrptb or xdrptb (8, 15).

We can envisage that models incorporating multiple validated CT features as well as clinical risk factors will have high reliability for predicting mdrptb.

## Acknowledgement

We thank Dr Xiao-Rong Wang and Dr Yu-Feng Ye, former visiting radiologists at the Chinese University of Hong Kong, for reading CT images with us. We also thank Ms Jing Wang at the Shenzhen Center for Chronic Disease Control, and Miss Liu Yang, former research assistant at the Chinese University of Hong Kong, for data collection at the Shenzhen site, China.

